# Stable Structures or poly(A)-binding protein loading protect cellular and viral RNAs against ISG20-mediated decay

**DOI:** 10.1101/2023.06.05.543696

**Authors:** Camille Louvat, Séverine Deymier, Xuan-Nhi Nguyen, Emmanuel Labaronne, Kodie Noy, Marie Cariou, Antoine Corbin, Mathieu Mateo, Emiliano P. Ricci, Francesca Fiorini, Andrea Cimarelli

## Abstract

ISG20 is an interferon-induced 3’-to-5’ RNA exonuclease that acts as a broad antiviral factor. At present, the features that expose RNA to ISG20 remain unclear, although recent studies have pointed to the modulatory role of epitranscriptomic modifications in the susceptibility of target RNAs to ISG20. These findings raise the question as to how cellular RNAs, on which these modifications are abundant, cope with ISG20. To obtain an unbiased perspective on this topic, we used RNAseq and biochemical assays to identify elements that regulate the behavior of RNAs against ISG20. The results we have obtained indicate that poly(A)-binding protein (PABP1) loading on the RNA 3’ tail provides a primal protection against ISG20, easily explaining the overall protection of cellular mRNAs observed by RNAseq. The second element we uncovered is provided by terminal stem-loop RNA structures, that have been associated to ISG20 protection before, but that we re-examine here systematically to define the stabilities that tilt the balance between resistance and susceptibility to ISG20. Given that these elements are present on cellular mRNAs, but can be co-opted by viruses as well, these results shed new light on the complex interplay that regulates the susceptibility of different classes of viruses against ISG20.

## INTRODUCTION

The interferon sensitive gene of 20kDa (ISG20) is an antiviral RNase that belongs to the DnaQ-like exonuclease superfamily, which is highly conserved across prokaryotes and eukaryotes. Members of this superfamily present a common catalytic core defined by four conserved aspartate and glutamate residues (DEDD) and can exhibit either RNase or DNase 3’-to-5’ exonuclease behaviors (1, 2). Among the members of this family, ISG20 was identified in 1997 as a type-I interferon (IFN)-induced protein and was soon associated to RNA virus inhibition (3, 4, 5, 6). Over the years, the spectrum of viruses described to be inhibited by ISG20 has expanded to encompass several positive-and negative-strand RNA viral families, in addition to Retroviruses (7,8, 9, 10, 11, 12, 13, 14, 15, 16, 17, 18, 19, 20, 21). However, a number of studies have also identified viruses that exhibit a strong resistance to this antiviral factor, pointing on the whole to the existence of a likely complex relationship between ISG20 and viruses that remains to be fully appreciated (11,16,17).

Similarly, the exact mechanism of viral inhibition by ISG20 remains unclear (6). In light of its strong RNA exonuclease activity, the first mechanism of viral inhibition proposed for ISG20 was based on the direct degradation of viral RNA (4,5). This model was supported by the fact that ISG20 behaves as a potent RNase *in vitro* and that lower levels of viral RNA were often, albeit not always, measured in ISG20-positive cells undergoing infection. Also in agreement with this model, point mutations in the catalytic core of ISG20 compromised both its ability to degrade RNA *in vitro* and its ability to inhibit viral replication in cells (4,5). However, several studies failed to report a strong decrease of viral RNA levels despite clearly measurable antiviral effects of ISG20, raising the possibility that alternative mechanisms, among which translation inhibition from viral RNAs, could be at play (11,16,17, 18).

A recent feature that gained interest as a possible explanation for how ISG20 could discriminate between RNAs has been the presence of epitranscriptomic modifications, such as N^6^-Methyladenosine (m^6^A), or 2’O-Methylation (2’O-Me) (19) (22). These and other modifications are added co-transcriptionally in the nucleus on cellular RNAs and can influence virtually every aspect of their metabolism, among which nuclear export, stability or translation rates. These modifications are dynamic, as they can be removed by dedicated enzymes referred to as erasers and can also act as docking sites for specific proteins called readers, overall accounting for the pleiotropic influence that epitranscriptomic modifications play on the behavior of RNAs (reviewed in (23). An increasing number of studies are depicting a complex canvas on how viruses themselves can co-opt such modifications for their own purposes, among which to mimic cellular RNAs (reviewed in (24). In this respect, the presence of a single m6A modification at a specific location on the hepatitis B virus (HBV) viral RNA has been recently described to promote ISG20-mediated RNA degradation, through the recruitment of a m6A reader protein YTH N^6^-Methyladenosine RNA binding protein F2 (YTHDF2)-ISG20 complex (19). However, an opposite behavior has been described in the case of 2’O-Me modifications that are scattered along the human immunodeficiency type 1 virus (HIV-1) 9 kilobases genome and that act as protective elements against ISG20-mediated degradation (22). The fact that epitranscriptomic modifications can modulate the susceptibility of RNAs to the action of ISG20 opens up to the unaddressed question of how this enzyme behaves towards cellular RNAs in which these modifications are largely present. To more globally appreciate the effects of ISG20 on RNAs at the scale of the whole cells, we performed an RNAseq analysis on cellular mRNAs. The results we have obtained indicate that ISG20 expression does not lead to overt changes in the cellular transcriptome suggesting that cellular mRNAs are generally protected against ISG20. However, a small effect was observed on histones mRNAs that represented the main category of mRNAs downregulated in the presence of ISG20. This finding is of interest because histone mRNAs are the only mRNA species in the cell devoid of poly(A) tails, but instead exhibit a 3’ stem-loop structure (25), which are surprisingly similar to those that are often described as protective against ISG20-mediated degradation (14). This prompted us to re-examine the reasons and the determinants that govern the resistance of poly(A) cellular mRNAs on one hand and the susceptibility of stem-loop bearing RNAs using finer biochemical analyses that used purified recombinant ISG20 protein and synthetic model RNAs. The results we have obtained indicate that the poly(A)-binding protein (PABP1) provides a key layer of protection against ISG20, by shielding the 3’ extremity of cellular mRNAs, providing on one hand the explanation for the resistance of cellular mRNAs from ISG20 and on the other opening the question of the interplay between ISG20 and the plethora of viral and cellular binding factors that often decorate RNAs. In the case of non-poly(A) mRNAs, we confirm that stem-loop structures can similarly protect RNA from ISG20 degradation, but only if their thermodynamic stability is equal or superior to 20 Gibbs free energy (ΔG).

Overall, this study identifies two key features that mediate RNA protection from ISG20, shedding novel light on the complexity of the relationship between ISG20 and its viral RNA targets.

## MATERIALS AND METHODS

### Cell culture and DNA constructs

Human kidney epithelial HEK293T and lung epithelial A549 cells (ATCC: CRL-3216 and CCL-185, respectively) were maintained in complete Dulbecco’s Modified Eagle Medium (DMEM, Gibco, cat. 61965-026) supplemented with 10 % fetal bovine serum (Sigma, cat. F7524) and 100U/mL of penicillin-streptomycin (Gibco, cat.11548876). Transient transfections were performed using an in-house Calcium-Phosphate (Calcium-HBS) method (26). Two series of plasmids coding for an N-terminal Flag-tagged ISG20 (*wild-type* and D94A mutant, referred to as WT and M1, respectively) were used, as previously described (17): pcDNA3.1+-based plasmids used for ectopic expression and pRetroX-tight-Puro-based ones (Clontech, cat. PT3960-5) used here to generate stable cell lines expressing ISG20 upon induction with doxycycline (dox.), following retroviral-mediated gene transduction (see below).

The following antibodies were used for Western Blotting: anti-Flag (Sigma cat. F3165) anti-α-Tubulin (Sigma cat. T5168), anti-GFP (Sigma cat. G1544), anti-mouse, and anti-rabbit IgG-Peroxidase conjugated (Sigma, cat. A9044 and cat. AP188P, respectively).

### Generation of stable cell lines

Murine leukemia virus (MLV) retroviral particles were obtained by transient transfection of HEK293T cells with plasmid DNAs coding for: the MLV structural protein Gag-Prol-Pol (pTG13077, (27); the envelope protein G from the Vesicular Stomatitis Virus (VSVg); and the mini-viral genome coded by the pRetroX-Tight-Puro (8:4:8 μg DNA per 10 cm plate dish, respectively). A parallel virus production was obtained by substitution of the ISG20 coding viral genome with pRetroX-Tet-On (Clontech, cat. 632104) that codes for the TetOn protein and allows for dox. inducible expression of ISG20. Transfection with these three plasmids allows the production of MLV-derived retroviral particles that can be used to obtain stable insertion of the transgene of interest into the genome of target cells. Viral particles released in the supernatant of transfected HEK293T cells were harvested twenty-four to forty-eight hours after transfection, syringe-filtered (0.45 μm) and used to challenge A549 cells. Following transduction with both types of particles (pRetroX-Tight-Puro-ISG20 and pRetroX-Tet-On), cells were selected as a pool upon incubation with Puromycin and G418 (Sigma, cat. P9620 and G418-RO). ISG20 expression was routinely induced upon incubation of cells for 18 hours with 1.5 μg/mL of doxycycline (Takara Bio, cat. 631311).

### Mopeia Virus infections

A549-ctl, -WT or -M1 cells were plated at 250,000 cells per well in 24-well plates in presence of doxycycline (1.5 μg/mL) twenty-four hours prior to infection with a recombinant MOPV-WT (strain AN21366, (28) at a multiplicity of infection (MOI) of 0.01. Supernatants and cells were collected forty-eight hours later for further analyses. Viral loads in cells and cell supernatants were measured by RT-qPCR. To this end, RNAs were extracted using a QIAamp Viral RNA kit (Qiagen, cat. 52904) and amplified using the SensiFAST Probe No-ROX One-Step kit (Bioline, cat. BIO-76001) using primers and probe targeting the NP gene of MOPV (**Supplementary Table 1**). Cellular RNAs were instead extracted using an RNAeasy mini extraction kit (Qiagen, cat. n. 74004) and then treated as described above. Antibodies were: rabbit anti-Z antibodies (produced in house), HRP-conjugated anti-FLAG (SIGMA, cat. A8592) and anti-Actin antibodies (SIGMA, cat. A3854).

### RNA-seq

HEK293T cells were seeded in 10-cm dishes and cells were transfected twenty-four hours later with 10 μg of pcDNA3.1+ control vector or of vector expressing Flag-ISG20 WT or M1, along with a vector coding for a GFP reporter that we used as an internal control. Twenty-four hours after transfection, total RNAs were extracted using Trizol added directly on the culture plate. Extracted RNAs were depleted from ribosomal RNAs using antisense DNA oligonucleotides complementary to rRNA and RNaseH as previously described (29), followed by DNase treatment to remove DNA oligonucleotides. High-throughput sequencing libraries were prepared as described (30). Briefly, RNA samples depleted from ribosomal RNAs were fragmented using RNA fragmentation reagent (Ambion, Cat: AM8740) for 3 minutes and 30 seconds at 70°C followed by inactivation with the provided “Stop” buffer. Fragmented RNAs were then dephosphorylated at their 3’ end using T4 Polynucleotide kinase (PNK, New England Biolabs, Cat: M0201) in MES buffer (100 mM MES-NaOH, pH 5.5, 10 mM MgCl2, 10 mM β-mercaptoethanol and 300 mM NaCl) at 37 °C for 3 h. RNA fragments with a 3′-OH were ligated to a preadenylated DNA adaptor. Following this, ligated RNAs were reverse transcribed with Superscript III (Invitrogen, cat. 18080044) with a barcoded reverse-transcription primer that anneals to the preadenylated adaptor. After reverse transcription, cDNAs were resolved in a denaturing gel (10% acrylamide and 8 M urea) for 1 hour and 45 minutes at 35 W. Gel-purified cDNAs were then circularized with CircLigase I (Lucigen, cat. CL4111K) and PCR-amplified with Illumina’s paired-end primers 1.0 and 2.0. PCR amplicons (12-14 cycles for RNA-seq and 4-6 cycles for ribosome profiling) were gel-purified and submitted for sequencing on the Illumina HiSeq 4000 platform.

### Analysis of high-throughput sequencing reads

Sequencing reads were split with respect to their 5′ in-line barcode sequence. After this, 5′-barcode, 6nt Unique Molecular Identifier (UMIs) were removed from reads using an in house script. Following this, the first 7 nucleotides at the 5’ end of R1 reads and the first 60nt at the 3’ end of R2 reads were further removed to avoid dovetails. 3’ adaptor sequences were removed using Trimmomatic (31) with the following parameters “PE MAXINFO:36:0.2”. To remove rRNA and tRNA contaminant sequences, reads were first mapped to a custom set of sequences including human 45S and 5S rRNA, tRNAs, phiX174 and GFP sequences using Bowtie2/2.3.3 (32) with the following parameters “bowtie2 -t --sensitive”.

Reads that failed to map to this custom set of sequences were next aligned to the human hg38 assembly and the GRC-Genecode GRCh38.v37 primary assembly annotation using HISAT2 2 (v2.1.0) (33) with the following parameters “hisat2 -t --no-unal --phred33 -p 16 -k 10 --min-intronlen 20 --max-intronlen 1000000 --rna-strandness RF I 100 -X 700 --fr”. Read counts on all transcripts of interest were obtained using the HTSeq count package (34) with the following parameters “htseq-count -f sam -r pos -s yes -a 10 --nonunique=none -m union”.

### Differential analysis with DESeq2

Differentially expressed genes upon ISG20 overexpression were obtained using DESeq2 (35) (version 1.24.0) in R (version 3.6.3).

### Proteins expression and purification

The *wild-type* ISG20 full coding sequence (amino acids 1–181) was cloned into the pPROEX HTa vector (Thermo Fisher Scientific, cat. 10711018) by standard molecular biology techniques. Protein expression from this plasmid results in the presence of a N-terminal hexa-histidine tag-ISG20 fusion. The pTRC-HisA plasmid harboring the PABP1-coding sequence was kindly provided by Dr. Théophile Ohlmann (CIRI, Lyon, France).

ISG20 was expressed in *E*.*coli* BL21(DE3) Rosetta/pLysS strain (Thermo Fisher Scientific, cat. EC0114 and Sigma Aldrich, cat. 71403, respectively). Bacterial cells were grown in an LB medium for 16 hours at 25°C, then protein expression was induced with 0.1 mM IPTG (isopropyl-β-d-thiogalactopyranoside, Sigma Aldrich, cat. PHG000110). Cultures were incubated 3h30 at 25°C under continuous shaking then harvested by centrifugation for 15 minutes at 5000 g. Each pellet was resuspended with buffer A (50 mM Tris-HCl pH 7.5, 150 mM NaCl, 20 mM imidazole, 5 mM β-mercaptoethanol, and 20% glycerol, v/v) supplemented with 1 mM of adenosine triphosphate (ATP), 1 mM MgCl2, and Protease Inhibitor Cocktail tablet (ROCHE, cOmplete™ Sigma Aldrich cat. 11836170001). The soluble lysate was applied to a prepacked nickel column (HisTrap HP column, Cytiva Europe GmbH, France cat. GE17-5247-01) and fractionated on an AKTA pure system (Cytiva Europe GmbH, France) using a linear gradient from buffer A to buffer B (buffer A supplemented with 0.5 M imidazole) over 10 column volumes. A second step of purification was carried out using a Superdex 75 10/300 GL column (Cytiva Europe GmbH, France) with an isocratic elution carried out with storing buffer (50 mM Tris-HCl pH 7.5, 50 mM NaCl, 2 mM β-mercaptoethanol, and 20% glycerol, v/v). Finally, glycerol was added to reach the concentration of 50% and proteins were stored at -20°C.

Bacterial cells for His-tagged PABP1 expression were grown in LB medium at 37°C before induce protein expression overnight at 20°C with 1 mM IPTG and shift the culture to 20°C. The pellet was resuspended in a lysis buffer composed of 50 mM Tris-HCl pH 7.5, 300 mM NaCl, 20 mM imidazole, 5 mM β-mercaptoethanol, and 20% glycerol and purification was performed as described above. A second step of purification was carried out using a Superdex 75 10/300 GL column with high ionic stringent buffer (50 mM Tris-HCl pH 7.5, 1 M NaCl, 2 mM β-mercaptoethanol, 0.1 mM EDTA and 20% glycerol). The protein was dialysed against storing buffer (50 mM Tris-HCl pH 7.5, 50 mM NaCl, 2 mM β-mercaptoethanol, and 20% glycerol, v/v) then stored at -80°C.

### *In vitro* RNA synthesis, purification, and radiolabeling

RNAs were produced following transcription from partially double-stranded DNA templates using the T7 RNA polymerase enzyme mix (Thermo Fisher Scientific cat. EP0112), following the manufacturer’s instructions and as described in (36). TAR (nucleotides 454-512 of the HIV-1 proviral genome # AF004394) and TAR-9SL DNA templates were produced by the annealing of two complementary oligonucleotides (**Supplementary Tables 1** and **2**). RNAs were dephosphorylated using Calf Intestinal Alkaline Phosphatase (New England Biolabs, cat. M0525) then purified on denaturing 10% (w/v) polyacrylamide gel (29:1) as previously described (36). Before use in binding and enzymatic studies, dsRNA was heated in a refolding buffer (20 mM HEPES pH 7.5, 0.2 M NaCl, 2 mM MgCl2, 2 mM DTT) for 3 minutes at 95 °C followed by 40 minutes of slow controlled cooling to room temperature and finally placed on ice. TAR-5SL and poly(A) RNAs were chemically synthesized (Dharmacon). Radiolabeling of 50 pmol of RNA substrate was performed in 20 μl of reaction mix using 10 U of T4 PNK (New England Biolabs, cat. M0201) and 3μL of ?^32^P-ATP (Perkin Elmer) in 1X PNK buffer. The mixture was incubated at 37°C for 1 h before inactivating the reaction by incubating at 65°C for 20 minutes. Radiolabeled RNA was first purified using Microspin G25 column (Sigma Aldrich cat. GE27-5325-01) following the manufacturer’s instructions, then extracted with an equal volume of Phenol:Chloroform:Isoamyl Alcohol mix (25:24:1; Sigma Aldrich cat. 516726-1SET) precipitated with ethanol.

### Electrophoretic Mobility Shift Assays (EMSA)

Samples were prepared by mixing a radiolabeled RNA (10 nM) with recombinant ISG20, as indicated in the text, or in the legend, in a buffer containing 50 mM Tris-HCl pH 7.4, 150 mM NaCl, 5 mM β-mercaptoethanol, 10 mM MnCl2, 0.1 μg/μL of BSA, and 5% (v/v) glycerol. The samples were incubated at 30°C for 30 minutes before being resolved by native 6% polyacrylamide (19:1) gel electrophoresis and analyzed by phosphorimaging. For PABP1 EMSA, 10 nM of Poly(A)-containing substrate was incubated with PABP1 (50, 100, 200 and 500 nM) in a buffer containing 25 mM Tris-Hcl pH 7.4, 5 mM MgCl2, 100 mM NaCl, 2.5 mM DTT, 0.2 μg/μL of BSA, and 10% (v/v) glycerol. Protein and RNA were incubated at 37°C for 15 minutes before being resolved by native gel, as previously indicated.

### 3’-5’ Exonuclease assays

Nuclease assay was performed by mixing 5 nM of recombinant ISG20 with 500 nM of radiolabeled RNA substrate in a buffer containing 50 mM Tris-HCl pH 7.4, 2.5 mM MnCl2, 1 mM β-mercaptoethanol, 0.4 mM DTT, 0.1% (v/v) Triton X-100, and 10% (v/v) glycerol. The reaction was incubated at 37°C for 1 hour. At the end of each incubation time 3 μl aliquots were withdrawn and rapidly mixed with 3 μL of stop buffer containing 5 mM EDTA, 0.5%(w/v) SDS, 34%(w/v) glycerol, 0,5 M urea, 1% (w/v) formamide, 0.01%(w/v) xylene cyanol and 0.01%(w/v) bromophenol blue. The collected samples were heated at 98°C for 5 minutes then resolved in 15% polyacrylamide (ratio 19:1) - 7 M urea gel and analyzed by phosphorimaging using Typhoon FLA 9500 (Cytiva Europe GmbH). When indicated the bands were quantified using ImageQuant software (Cytiva Europe GmbH) and the results analyzed by KaleidaGraph (Synergy software). Processivity tests were performed using cold RNA to trap the excess enzyme. ISG20 was previously incubated with radiolabeled RNA substrate for 5 minutes then 100 molar excess of cold RNA substrate was added when indicated. Exonuclease reactions were subsequently performed as described above.

### PABP1-protection exonuclease assays

Thirty nM of RNA substrate was mixed with 900 nM of recombinant PABP1 and incubated for 20 minutes at 37°C in a buffer containing (10 mM Tris-HCl pH 7.4, 10 mM MnCl2, 1 mM DTT, 100 mM NaCl, and 10% glycerol). ISG20 was added at final concentration of 120 nM and the enzymatic assay was conducted as described above.

### Statistical analyses

Statistical analyses were calculated with the Graphpad Prism8, or Excel softwares: Student t tests (unpaired, two-tailed), or one-way Anova with Dunnett’s multiple comparison tests, as indicated in the legend of the relevant figures.

## RESULTS

### ISG20 does not drive major changes in the cellular transcriptome, but leads to a small decrease in histone mRNAs

To determine the effects of ISG20 on cellular mRNAs at the whole cell scale, the DNA coding ISG20 proteins was ectopically transfected in HEK293T cells, prior to WB and RNAseq analyses twenty-four hours later (**Figure 1A** and **1B** for WB). WT ISG20 was compared to an ISG20 mutant presenting a single point mutation (M1, or D94A) within the conserved DEDD residues quartet that abrogates the ability of ISG20 to degrade RNA (**Figure 1B**) (17).

**Figure 1.**
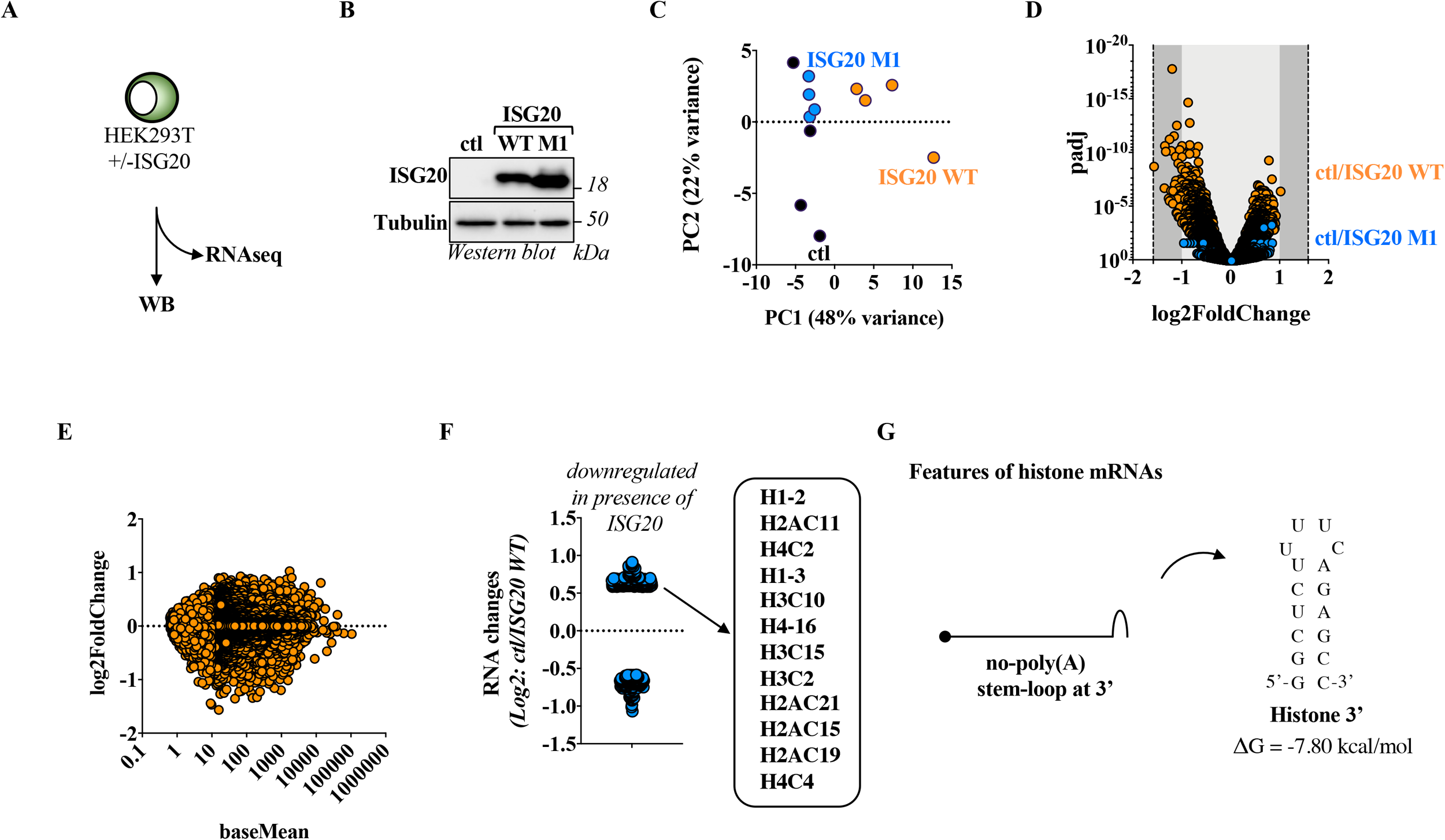
ISG20 does not lead to overt changes in the cellular mRNA transcriptome with the exception of a small effect on histone mRNAs. A) HEK293T cells were ectopically transfected with DNAs coding for control, WT or M1 (D94A) ISG20 proteins and twenty-four hours later, cells were lysed and samples were analyzed by WB (B) or used after ribosomal RNA depletion for RNAseq analysis (C to F). C) PCA analyses of the three conditions examined here. D) mRNA changes in the control versus WT or M1 ISG20 conditions (*abscissa*) plotted in function of the adjusted p-value (padj, *ordinate*). E) Analysis of the changes in individual genes (*ordinate)* as a function to the gene length (baseMean, *abscissa*). F) Highlight of genes significantly modulated by +/-1.5-fold in the presence of WT ISG20. G) Schematic representation of histone mRNAs and stem-loop stability prediction using the UNAFold web server (Zuker 2013 <u>10.1093/nar/gkg595</u>). The WB panels depict typical results obtained, while the graphs presented in the remaining panels presents data obtained from 4 independent samples.

RNA-seq analysis indicated a clear separation of the samples by principal component analysis (PCA, **Figure 1C**), however statistically significant changes in mRNA levels were contained within 2-fold and this independently from the levels of expression of a given cellular RNA (**Figure 1D** and **1E**), indicating that, under the experimental conditions used here, ISG20 does not lead to major changes in the cellular transcriptome. However, when the threshold was lowered to a 1.5 fold change, respectively 208 and 230 genes appeared as up and down-regulated in the presence of WT ISG20 (**Figure 1F**). Interrogation of the Reactome database (37) with the genes downregulated, and thus potentially targeted, by ISG20 indicated an enrichment in histone-related pathways (**Supplementary Figure 1**), essentially driven by changes in the levels of twelve distinct histone-coding mRNAs (**Figure 1F**). Histone-coding mRNAs constitute the sole example of non-poly adenylated mRNA, amongst cellular mRNAs. However, the 3’ end of histone mRNAs presents a common stem-loop structure (**Figure 1G** and **Supplementary Table 3**) (25), that are generally believed to protect against ISG20-mediated degradation. With the caveat that this RNAseq analysis alone does not prove that ISG20 directly degrades histone-mRNAs, these results opened up to two distinct but complementary questions: first, how the majority of poly(A) cellular mRNAs could be protected by the action of ISG20 and second, why histone mRNAs that present a 3’ terminal stem-loop that should protect them from ISG20, appear susceptible to this RNA exonuclease.

### The Mopeia Arenavirus is resistant to ISG20-mediated inhibition

To re-examine the possibility that stem-loop structures could indeed protect from ISG20 from a distinct angle, we studied the behavior of the Mopeia virus (MOPV) in A549 lung epithelial cells expressing a dox.-inducible form of ISG20. MOPV belongs to the *Arenaviridae* family and is a bisegmented ambisense RNA virus. The segmented portions of the genome of *Arenaviridae* code for genes in opposite polarity that converge into a common genome portion named intergenic region (IGR). This region is characterized by long palindromic stretches that fold into highly stable stem-loop structures (**Supplementary Table 4**) that are thus present at the 3’ of all viral RNAs (**Figure 2A**, the plus strand orientation of the IGR sequences of the long and short segments of the MOPV genome, L and S respectively, are presented) (38) (39).

**Figure 2.**
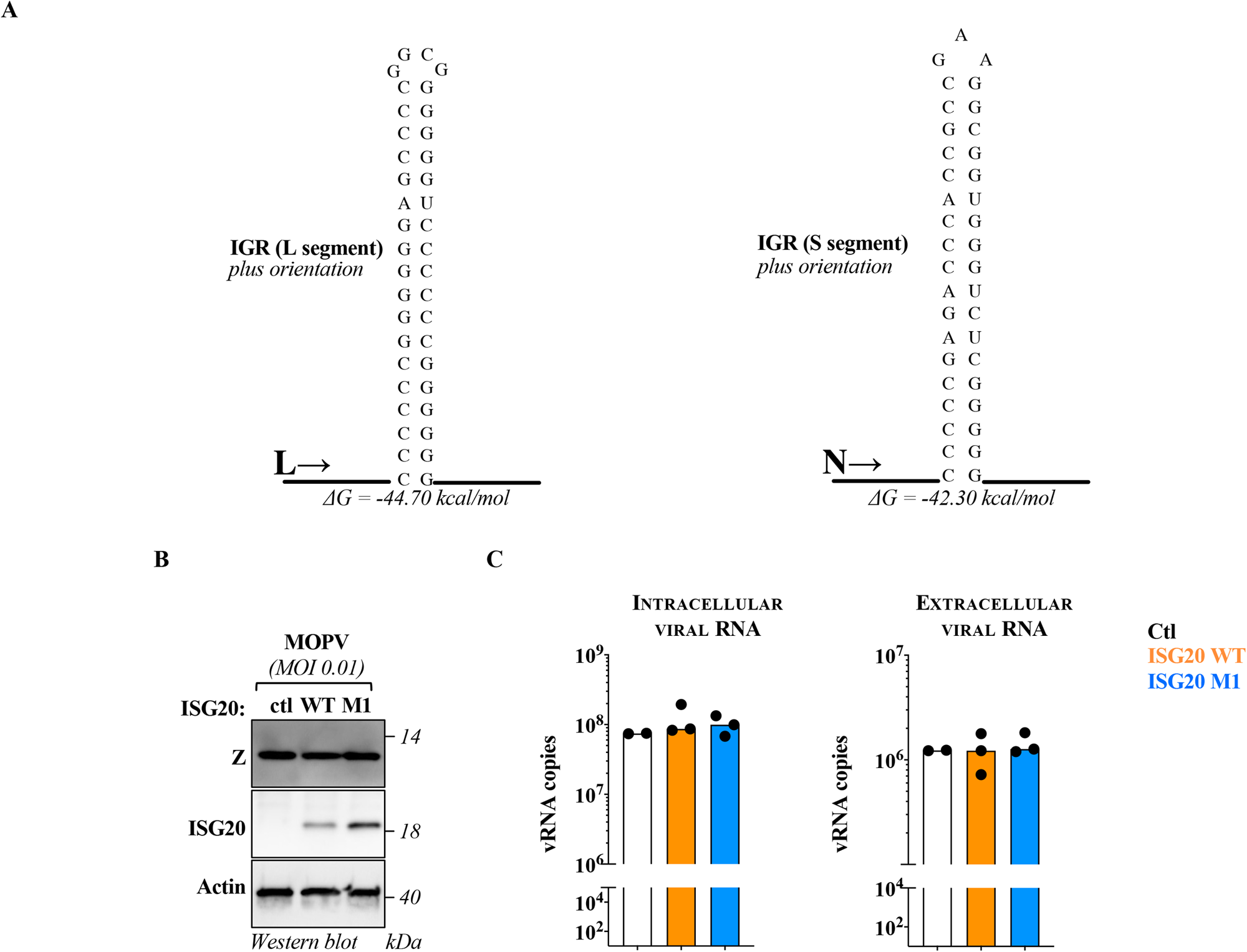
The Arenavirus Mopeia is resistant to ISG20-mediated inhibition. A) RNA structures of the intergenic regions (IGR) of the L and S segment of Mopeia virus, as determined by RNA Fold. B and C) A549 cells stably expressing WT or M1 ISG20 proteins were challenged with an MOI of 0.01 of MOPV and both cells and supernatants were analyzed forty-eight hours later for WB and RT-qPCR analyses. The WB panel presents typical results obtained, while the graphs present data obtained from 3 independent experiments carried out in duplicate (2 for control). Differences between control and remaining samples were not statistically significant following a one-way Anova with Dunnett’s multiple comparison test.

A549 cells expressing WT or M1 (D94A) ISG20 proteins were challenged with an MOI of 0.01 of MOPV and both cells and supernatants were analyzed forty-eight hours later to measure the accumulation of viral proteins by WB or the accumulation of viral RNA in the cell as well as in virion particles released in the cell supernatant (**Figure 2B** and **2C**). Under these conditions, ISG20 did not affect either the intracellular or extracellular viral RNAs quantities, indicating that MOPV is indeed resistant to ISG20 and strongly supporting the notion that structured 3’ ends can indeed act as an element of RNA resistance of viruses to ISG20 (**Figure 2C**).

### ISG20 acts as a distributive enzyme

The behavior of viral or cellular RNAs depends on a plethora of factors that can overall influence the susceptibility to ISG20 and also potentially explain the results mentioned above. It appeared then of importance to turn to reductionist methods to identify and study the reasons for the behavior of cellular and viral RNAs towards ISG20.

To explore this question, we purified ISG20 (harboring a hexahistidine N-terminal tag and expressed/purified from a prokaryotic expression system, **Supplementary Figure 2)** and characterized its behavior in biochemical assays. The first question we decided to address was whether ISG20 utilized a processive, or a distributive mechanism for its exonuclease activity. This is a fundamental issue to apprehend the mechanism of action of ISG20, but this enzymatic property has never been properly analyzed. To this end, we conducted a steady-state processivity assay (**Figure 3A**) by incubating purified ISG20 with radiolabeled linear ssRNA (called “hot”, 40 nucleotides long) in the presence, or absence, of an excess of cold ssRNA competitor in a reaction buffer containing Mn^2+^ as detailed before. The addition of ISG20 triggered the exonucleolytic reaction and the products of the reaction were monitored over 30 min by acryl-urea PAGE gel analysis (**Figure 3B**). As expected, ISG20 degraded completely its RNA substrate over the thirty minutes of the assay. However under the experimental conditions used here, ISG20-mediated degradation was completely prevented in the presence of the cold RNA competitor, indicating that ISG20 degrades RNA according to a distributive mode of action.

**Figure 3.**
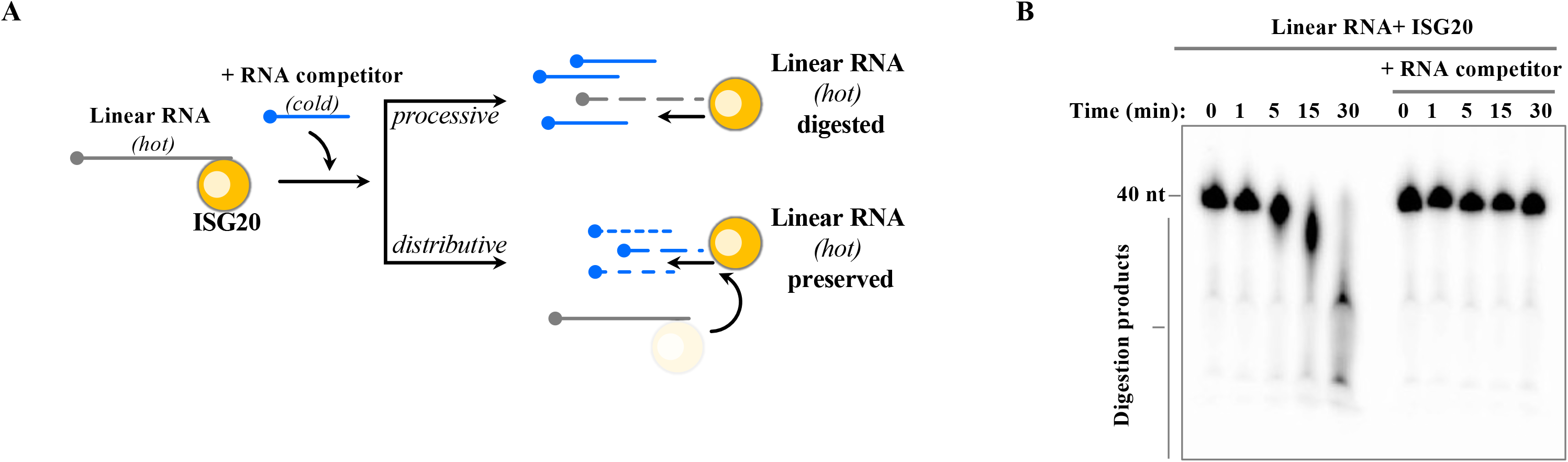
ISG20 behaves as a distributive enzyme *in vitro*. A) Schematic representation of the assay used to determine whether ISG20 acts as a processive or distributive enzyme. B) Linear target RNA was incubated with ISG20 in the presence or absence of cold ssRNA and the reaction was arrested at the indicated time prior to acryl-urea PAGE gel analysis of ISG20-mediated digestion. The panel presents typical results out of three independent experiments.

### ISG20-mediated degradation of a poly(A)-containing RNA target is prevented by PABP1

Given that expression of ISG20 in cells does not result in overt cellular RNA degradation and that ISG20 is a distributive enzyme, we hypothesized that a *trans*-acting factor such as PABP1 could shield poly(A) cellular mRNAs from ISG20-mediated degradation.

First, we expressed and purified recombinant PABP1 (**Supplementary Figure 2**) then verified its binding on a 5’ radiolabeled linear RNA containing 20 adenosines at its 3’ end by Electrophoretic Mobility Shift Assay (EMSA, **Figure 4A and 4B**). As expected, a PABP1:RNA complex was formed in a manner proportional to the concentration of PABP1. Next, we evaluated the possibility that PABP1 could interfere with ISG20-mediated degradation by shielding the poly(A)-tail by performing a steady-state exonuclease assay in the presence of non-saturating concentration of the PABP1 (**Figure 4C**). The substrate was first incubated for 20 minutes with PABP1, then ISG20 was added and its activity was monitored over a one-hour period. Under these conditions, a strong protection was exerted by PABP1 against the ISG20-mediated RNA degradation. In contrast, ISG20 remained able to degrade its RNA target in control conditions or in the presence of BSA (**Figure 4C and 4D**). Overall, this data indicates that PABP1 is able to shield the 3’ extremities of RNAs from ISG20-mediated degradation. Given that poly(A) tails are mandatorily present on cellular mRNAs, this likely represents the key feature of protection of cellular RNA from ISG20.

**Figure 4.**
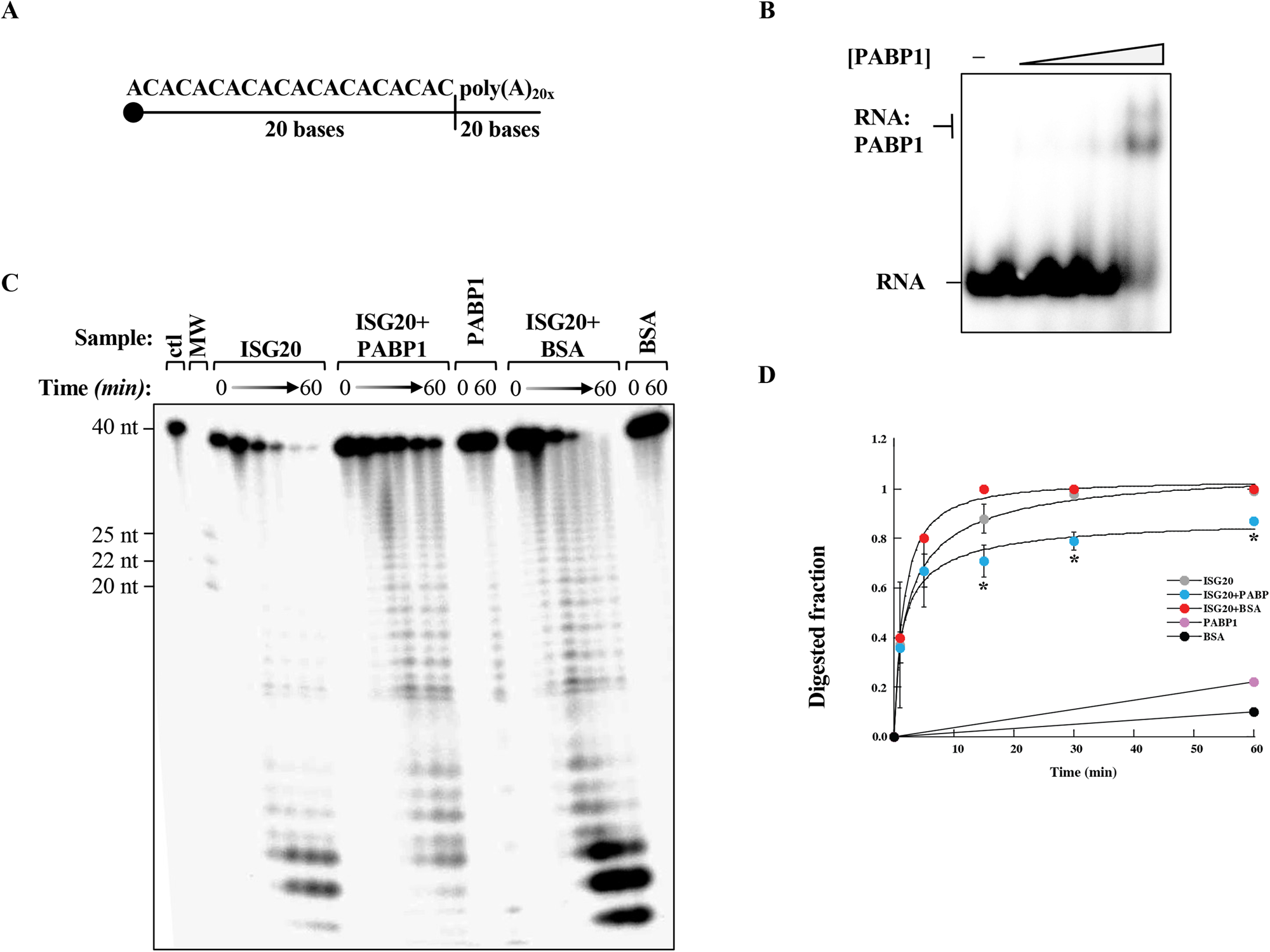
PABP1 protects poly-A tailed RNA from ISG20-mediated degradation. A) ssRNA sequence used in the protein protection assay bearing a poly(A) 3’ tail of 20 nucleotides. B) Representative EMSA illustrating the interaction of PABP1 with ^32^P 5’ end-labelled ssRNA. The RNA substrate (10 nM) was incubated with increasing concentrations of PABP1 (50, 100, 200 and 500 nM) or without protein, under the conditions described in Materials and Methods. C) Acryl-urea PAGE analysis of the exonuclease activity of ISG20 on the indicated ^32^P-labeled ssRNA either alone or pre-incubated with PABP1, or BSA. (D) Quantification of the protein protection assay presented in C. Graph showing the fractions of RNA digested by ISG20 as a function of time. The corresponding values for the digested fraction at time t (Dft) are calculated by the equation Dft= 1-(Rft/Rf0) where Rft is the intensity of the band corresponding to undigested RNA at time t and R0 at time 0, before the adding of the enzyme. Data derived from three independent experiments (mean ± SD). The best fitting with equation was found with Kaleidagraph (Synergy software) program. *, p<0.05 according to a Student t tests (unpaired, two-tailed) between the conditions ISG20 alone vs ISG20+PABP1 at the indicated time points.

### 3’ stem-loops protect RNAs from ISG20-mediated degradation, depending on their stability

The results obtained on the apparent higher susceptibility of histone mRNAs to ISG20 and the converse resistance of MOPV appear contradictory. However, the inspection of the thermodynamic stability of the stem-loop present on histone versus MOPV RNAs reveal important differences that could themselves be at the basis of their different behavior.

To explore this more formally and at a more general scale, we compared three well-characterized stem-loop structures derived from the transactivating region of the HIV-1 genome (TAR) and specifically the full TAR structure (TAR, spanning nucleotides 454-512 of the HIV-1 provirus (40)), along with two modified versions, TAR-9SL and TAR-5SL, in which the size of the stem has been reduced to 9 and 5 base pairs (**Figure 5A** and **Supplementary Table 5**; adapted from UNAFold; (41)). These structures were chosen because the folding thermodynamics computationally calculated at -32.90, -23.00 and -9.80, kcal/mol, respectively cover the spectrum of stem-loop stability that can be measured in histone, as well as MOPV mRNAs, thereby allowing us to define the boundaries of thermodynamic stability with which stem-loops can provide a sufficient protection against ISG20-mediated degradation.

**Figure 5.**
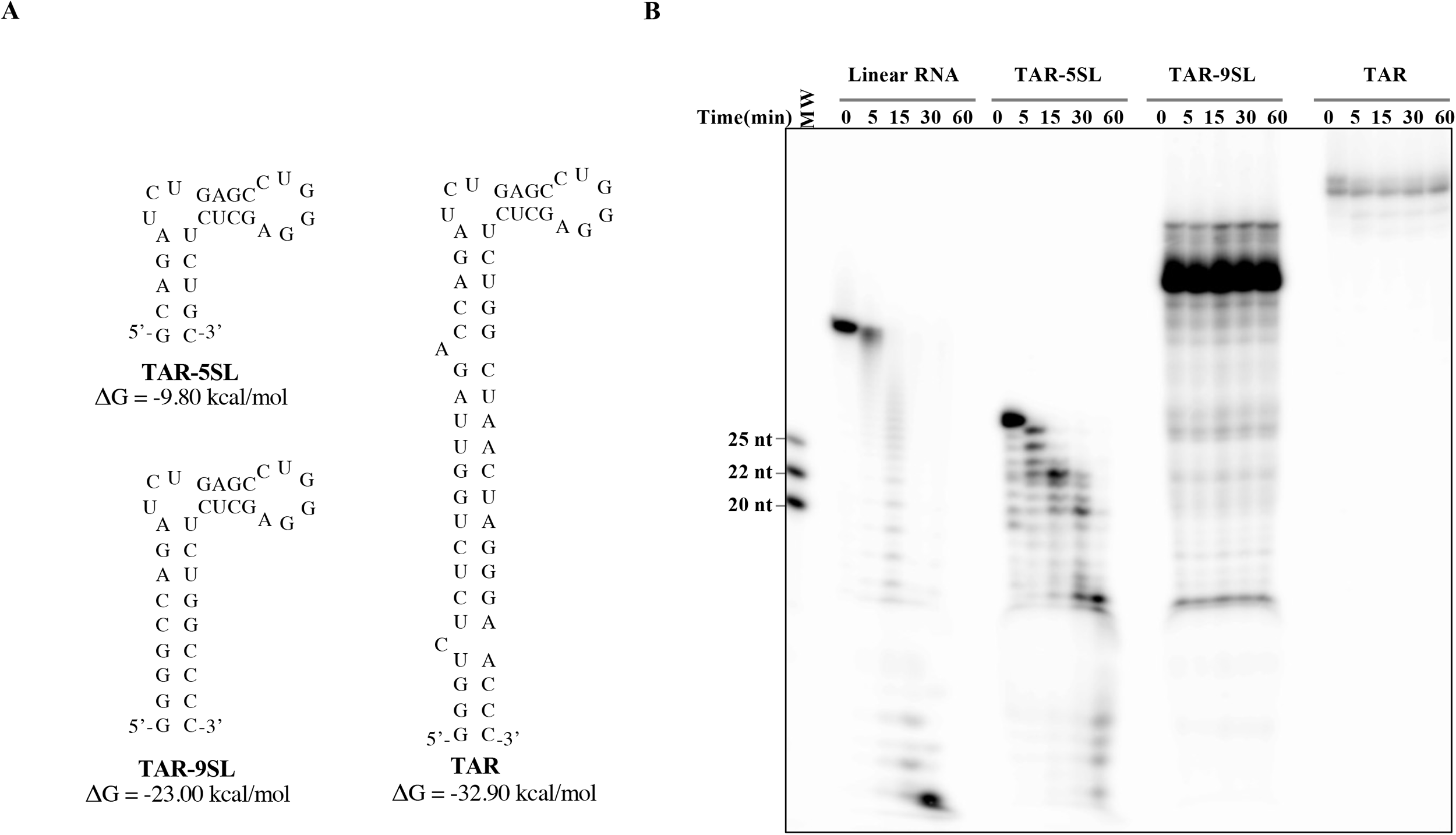
Only high energy structured 3’ RNAs provide protection from ISG20. A) Structural model of dsRNAs used here that differ for the length of their stem portions predicted with UNAFold web server (41) and Supplementary Table 5. B) Time course of ISG20-mediated digestion of linear single-stranded RNA (ssRNA, ie (AC)10A20), and the indicated stem-loop structures. The exonuclease reaction was analyzed by acryl-urea PAGE under the conditions described before. The gel depicts typical results obtained out of three independent experiments.

The different RNA substrates were radiolabeled, folded and purified, prior to use in an ISG20-mediated steady-state exonucleolytic reaction, as previously described in the Material and Methods section. We observed a lower intensity of labeling of the TAR structure compared to the others, which was not investigated further, but that we believe due to lower efficiency of T4 polynucleotide kinase activity on this substrate. As expected, the linear RNA substrate was degraded to completion very rapidly (**Figure 5B**). In contrast, the most stably structured RNAs (TAR-9SL and TAR) were completely resistant to ISG20 cleavage (**Figure 5B**). Instead, the TAR-5SL RNA presented a more complex pattern of digestion. In particular, a first ladder of five nucleotides corresponding to the degradation of the basal stem of the RNA appeared after five minutes of reaction, followed by the subsequent digestion of the 4-bp long apical stem and the complete digestion of target RNA at the end of the reaction. As such, these results indicate that stem-loop structures must possess a minimal stability to provide full protection against ISG20, while, short stem-loop remain susceptible to it. Of note, the lack of digestion of stable stem-loops structures is unlikely to be due to the absence of binding to ISG20, because ISG20 was able to associate to TAR-9SL in a classical EMSA assay (**Supplementary Figure 3**), suggesting that ISG20 bears the capability to sample RNAs even if they are highly structured.

## DISCUSSION

In this paper, we aimed to provide a more complete view of the mechanism(s) at the basis of the susceptibility/resistance of RNAs to ISG20-mediated degradation and to this end, we combined whole-cell RNAseq approaches to a biochemical characterization of the behavior of ISG20 against model RNAs. In so doing, we identified two novel elements of general protection of RNA from the degradative action of ISG20. The first is provided by PABP1, protein that associates with the poly(A)-tail present in cellular mRNAs (42). Under our experimental conditions and in line with the data obtained in cells, PABP1 binding to the 3’ ends of cellular mRNAs efficiently shields them from ISG20-mediated degradation, thus providing a simple explanation of why cellular RNA are essentially spared when ISG20 comes into action. The second element that our study identified, in line with previous results in the literature (4,14), is constituted by stem-loop structures and more precisely double-stranded regions that can protect RNAs by sequestering free 3’ ends from ISG20, even in the absence of a poly(A)-tail and its associated proteins. In this case however, we determined that the structure needs a minimal thermodynamic stability (≤ -23kcal/mol) to constitute an effective barrier against ISG20. Our results thus contribute to provide a general picture of elements that govern the behavior of ISG20 against RNAs.

A well-known example of antiviral RNAse in the cell is provided by the 2’-5’ oligoadenylate-dependent ribonuclease L (RNaseL), an endoribonuclease induced by viral infection, or type-I interferon responses (IFN-I) (reviewed in 43). The expression of RNaseL is tightly controlled at the transcriptional level, but also at a post-transcriptional one. Indeed, this enzyme is produced in an inactive form in the cell and it becomes activated in the presence of 5′-phosphorylated, 2′,5′-linked oligoadenylates, themselves produced by the 2′-5′ oligoadenylate synthase (OAS), another ISG whose activity is itself stimulated by double-stranded RNA (dsRNA). As such the activity of RNaseL is submitted to a multilayered and very tight mechanism of control that reflects its potential lethality for the cell (43). While ISG20 is also an ISG and is thus controlled at the transcriptional level, this enzyme does not seem to be produced in an inactive form, because ISG20 directly purified from cellular lysates behaves as a highly efficient RNase (17). While it remains possible that protein-cofactors lost during the purification procedure may thwart the RNase activity of ISG20 in the cell, whether such cofactors exist and how the cell restrains the activities of ISG20 remains unclear.

One possibility is therefore that ISG20 is able to discriminate RNAs according to features that remain to be identified. Recent studies have connected the presence of specific epitranscriptomic modifications in the modulation of the susceptibility of target RNAs to ISG20. Specifically, 2’-O nucleotide modifications have been clearly shown to protect the HIV-1 RNA genome from ISG20-mediated degradation according to a mechanism in which this bulky modification prevents the access of RNA to the catalytic site of ISG20 by steric hindrance (22). Conversely, a distinct epitranscriptomic modification (m6A) has been shown to expose HBV RNA to ISG20 digestion, by acting as a docking site for a protein complex composed by YTHDF2 and ISG20 that thus results in the direct deposition of ISG20 on its viral RNA target (19).

These findings are of interest because they open up to the more general question of how viruses and cells can exploit the large panel of epitranscriptomic modifications to influence most aspects of the RNA biology, including susceptibility to, or protection from, ISG20 that is an antiviral RNase. It remains for the moment unclear however, how epitranscriptomic modifications alone can provide a discriminating signal for ISG20. Indeed, these modifications are not only abundant on cellular RNAs, but also on RNAs from several classes of viruses, as an increasing number of studies is highlighting (44). Such modifications can be used for different purposes in the case of viruses, among which, to mimic cellular RNA and protect themselves from innate antiviral responses, or from antiviral effectors such as ISG20. An additional issue that remains to be addressed is how epitranscriptomic modifications, that often occur at internal positions of the RNA, can act distally to protect the 3’ end of target RNAs from ISG20. As such, the role of distinct epitranscriptomic modifications with respect to ISG20 is likely to be complex, multifactorial and perhaps context-dependent.

Our RNAseq data coupled with the fact that ISG20 acts distributively and that its activity is inhibited in the presence of PABP1, indicates that coating of the 3’ poly(A) by PABP1 provides an important and large spectrum element through which cellular mRNAs are protected from ISG20-mediated degradation. While it is true that certain classes of viruses also give rise to poly(A) RNAs, a large number of viruses does not, so that ISG20 may have evolved specifically to target them.

The spectrum of viruses that has been reported in the literature to be inhibited by ISG20 cannot be simply reconciled with the mere presence/absence of poly(A) tails, first because it is not known whether viral poly(A)-tailed mRNAs are always coated with PABP1 or with viral proteins and then because certain viruses exhibit 3’ extremities that are highly structured(7, 8, 9, 10, 11, 12, 13, 14, 15, 16, 17, 18, 19, 20, 21). In this respect, the subsequent finding of our study is that non-poly(A) tailed mRNAs can be protected by structured ends against ISG20. Several studies in the literature had reported that double-stranded RNAs and certain stem-loop RNA structures were resistant to ISG20-mediated degradation *in vitro* (17, 4,14). Starting from the observation that the major class of cellular mRNAs modified, albeit slightly, in the presence of ISG20 was represented by histone mRNAs and that these RNAs represent the unique example of mRNAs that is non poly(A)-tailed among cellular mRNAs, we carried out a more systematic analysis of stem-loop RNA thermodynamic stability in ISG20-mediated degradation and we compared stem-loop structures that spanned the differences in stabilities measured between histone mRNAs and MOPV RNAs. Our results indicate that ISG20 is able to sample complex stem-loops by binding to them. However, this sampling does not lead to target RNA degradation and in light of the distributive mechanism of action of ISG20, the enzyme is likely to come off its substrate afterwards. Instead, ISG20 efficiently degrades less stable stem-loops and our analysis indicates a thermodynamic stability comprised between -9.8 and -23 kcal/mol, as the boundary between susceptibility and resistance to ISG20.

These values ought to be considered as a first simplified reading of the action of ISG20, because in cells, these structures may be themselves regulated by the dynamic association of cellular or viral proteins, thus potentially influencing the extent and perhaps timing at which a given RNA can be susceptible to the action of ISG20. In the case of the stem-loop presents on histone mRNAs, this structure is normally associated with the Stem-Loop Binding protein (SLBP, 25) and it is thus possible that masking of this structure by a cellular protein can explain the low, albeit measurable, downregulation of histone mRNAs by ISG20.

Overall, our study uncovers two additional elements that can protect target RNAs from the degradative action of ISG20: poly(A)-tails and their default associated protein PABP1 and stem-loop structures, the minimal thermodynamic stability of which we define here. On more general grounds, our findings highlight the fact that the activity of ISG20 on target RNAs is likely modulated by a combination of factors among which the plethora of cellular or viral proteins that at any given time associate to target RNAs. These binding may not only protect RNAs from ISG20, by shielding the 3’ RNA ends, but on the other hand can also expose to it, for instance by unwinding specific stem-loop structures, overall shedding new light on the complexity that likely underlines the balance between susceptibility and resistance of RNA to ISG20.

## DATA AVAILABILITY

All data are incorporated into the article and its online supplementary material. RNAseq data is available at the Gene Expression Omnibus database of the NCBI as GEO Submission number GSE233792.

## SUPPLEMENTARY DATA STATEMENT

Supplementary Data are available at NAR online.

## ACKNOWLEDGEMENTS

We are grateful to Sylvain Baize for providing MOPV reagents.

## FUNDING

This work was supported by the Agence Nationale de Recherche (ANR, “grant number ANR-20-CE15-0025-01 to A.C., E. P. R. and S.D.). FF is supported by the CNRS (French National Centre for Scientific Research) and by FINOVI Foundation (Contract number 247479).

Funding for open access charge: Agence Nationale de Recherche

## CONFLICT OF INTEREST DISCLOSURE

The authors declare no conflict of interest.

## Notes

### Competing Interest Statement

The authors have declared no competing interest.

