## Supplemental Tables 1-5 and Supp Figures for "Stable Structures or poly(A)-binding protein loading protect cellular and viral RNAs against ISG20-mediated decay"

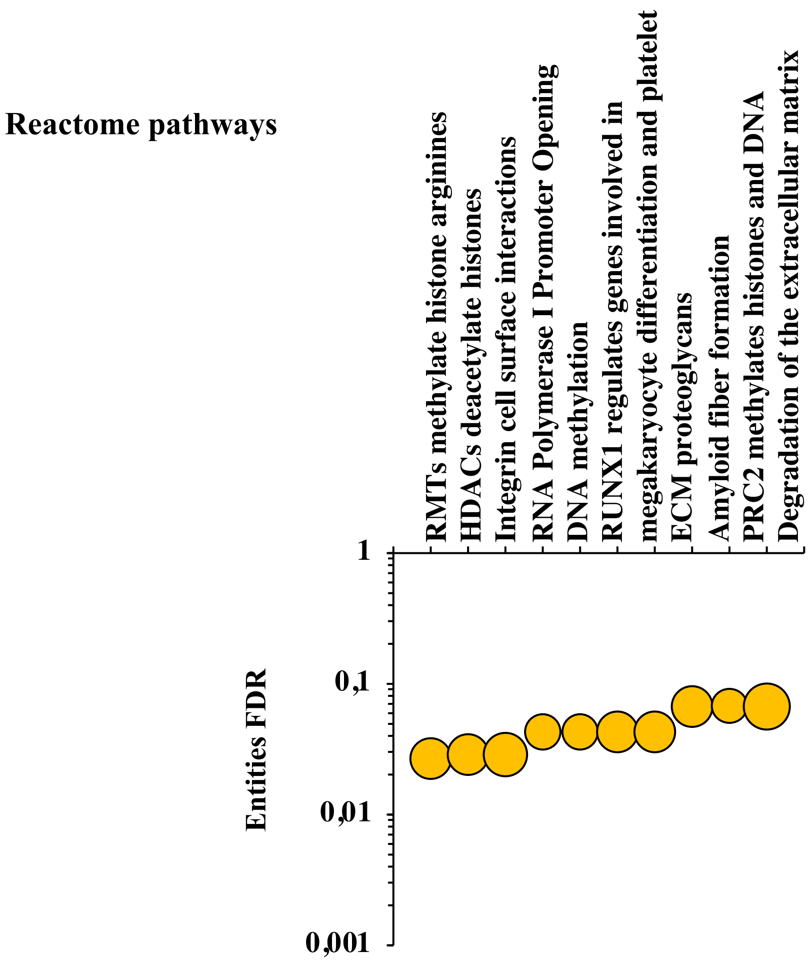


**Supplementary Figure 1.** Reactome pathway analysis of genes downregulated in the presence of WT ISG20. Only the top 10 pathways are shown here.

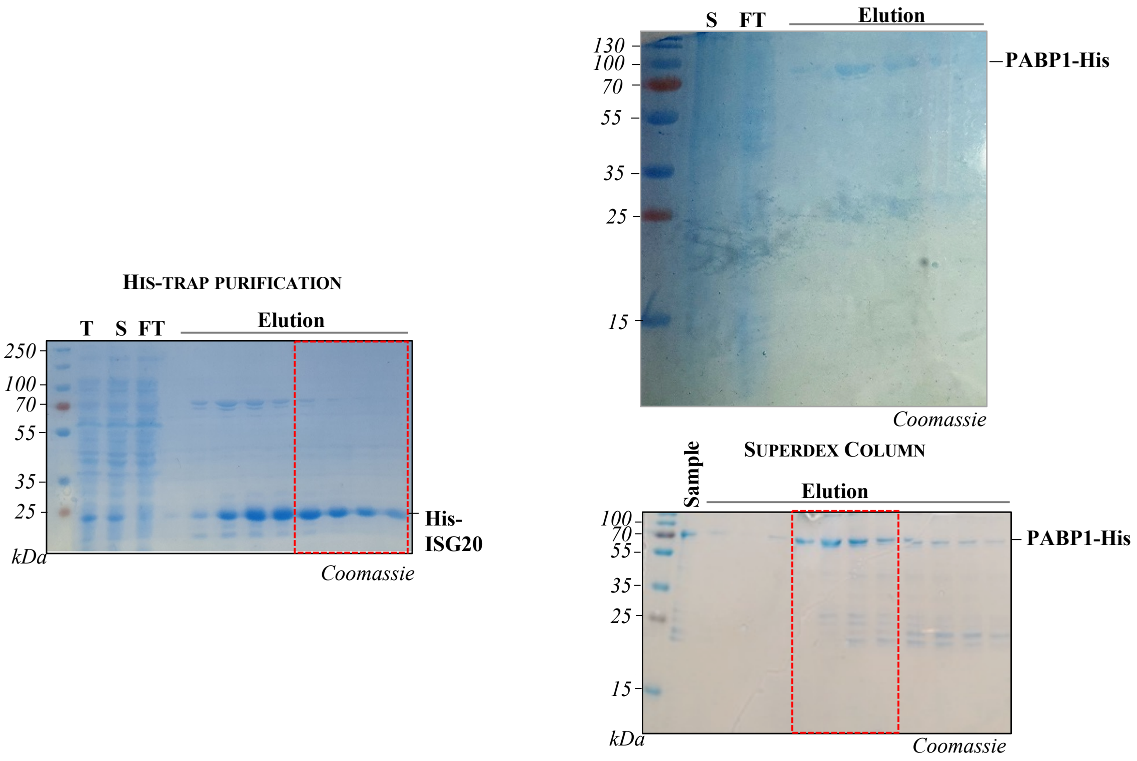


**Supplementary Figure 2.** SDS-PAGE gels illustrating the purification of his-tagged ISG20 and PABP1 expressed in *E. coli*, as indicated. Samples from the total lysate (lane T) and supernatant (lane S) fractions obtained after centrifugation of the E.coli lysate were loaded together with samples of the flow through (FT) and elution fractions obtained after nickel affinity chromatography or size-exclusion chromatography (Superdex Column, as indicated).

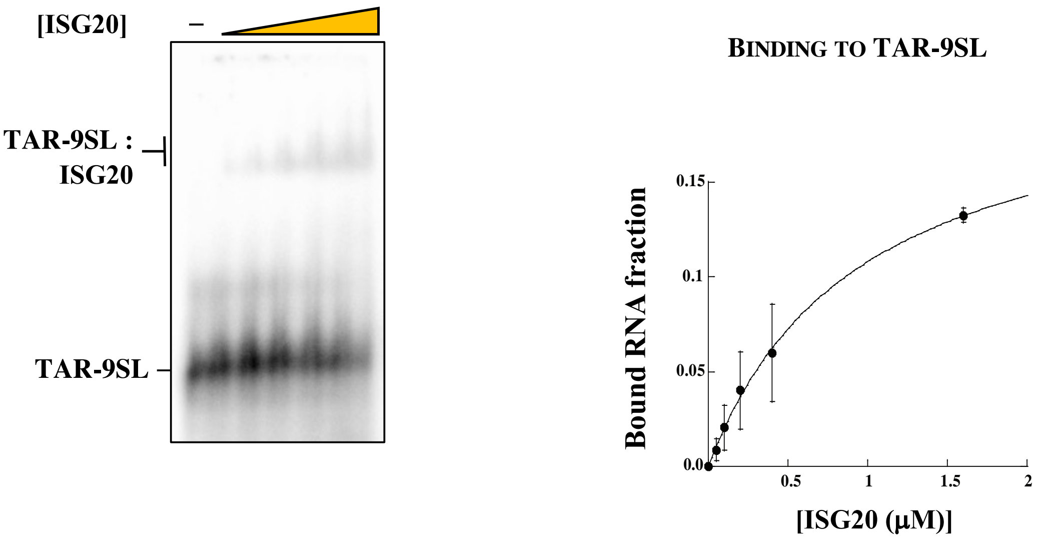


**Supplementary Figure 3.** Left) EMSA assay showing the interaction between ISG20 and the indicated structured RNA (TAR-9SL). The RNA substrate (10 nM) was incubated with increasing concentrations of ISG20 (50, 100, 200, 400 and 1600 nM) or without protein, under the conditions described in 'Materials and Methods'. Right) Quantification of ISG20 binding to dsRNA shown in figure A. The graph shows the fractions of RNA bound by ISG20 as a function of protein concentration. Detection and quantification of the bands were carried out using a Typhoon phosphorimaging system and the ImageQuant software (GE Healthcare). The bound product fraction at time 0 is calculated using the equation F0=P0/(R0+P0) and the product fractions at time "t" using Ft=(Pt-F0(Pt+Rt))/((1-F0)(Pt+Rt)) where P is the intensity of the shifted band and R the intensity of the substrate (RNA alone) in the corresponding EMSA gel. Data points (mean ± SD) derived from three independent experiments. Data were fitted with KaleidaGraph (Synergy software) using the Michaelis-Menten equation.

**Supplementary Table 1. Oligonucleotides used in this study.**

| Name | Sequence (5'- 3') |
|---|---|
| PT7 | AAATAATACGACTCACTATAGGG |
| FF3 | AAATAATACGACTCACTATAGGGGCCAGATCTGAGCCTGGGAGCTCTCTGGCCCC |
| FF4 | GGGCCAGAGAGCTCCCAGGCTCAGATCTGGTCCCTATAGTGAGTCGTATTATTT |
| FF5 | GGGTTCCCTAGTTAGCCAGAGAGCTCCCAGGCTCAGATCTGGTCTAACCAGAGAGACCCTATAGTGAGTCGTATTATTT |
| MOPV-NP-forward | GTCAAGCGTTCTTTGGGAATG |
| MOPV-NP-reverse | TCCAGAAAGACATAGTTTGTAGAGG |
| MOPV-NP-probe | FAM-TTCCTTTCCCCTGGCGTGTCA-BHQ1 |

**Supplementary Table 2. RNA substrates used in this study**

| Name | Sequence (5'- 3') | Synthesis |
|---|---|---|
| $(AC)_{10}A_{20}$ | ACACACACACACACACACACAAAAAAAAAAAAAAAAAAAA | chemically synthesized |
| TAR-5SL | GCAGAUCUGAGCCUGGGAGCUCUCUGC | chemically synthesized |
| TAR-9SL | GGGGCCAGAUCUGAGCCUGGGAGCUCUCUGGCCC | FF3/FF4 hybridization |
| TAR | GGGUCUCUCUGGUUAGACCAGAUCUGAGCCUGGGAGCUCUCUGGCUAACUAGGGAACCC | PT7/FF5 hybridization |

**Supplementary Table 3 mFold analysis of Histone 3' stem-loop structure.**

| **Histone structural element** | **δG** | **Information** |
|---|---|---|
| External loop | 0.00 | 0 ss bases & 1 closing helix |
| Stack | -3.30 | External closing pair is G1-C16 |
| Stack | -3.40 | External closing pair is G2-C15 |
| Stack | -2.10 | External closing pair is C3-G14 |
| Stack | -2.40 | External closing pair is U4-A13 |
| Stack | -2.10 | External closing pair is C5-G12 |
| **Helix** | -13.30 | 6 base pairs |
| Hairpin loop | 5.50 | Closing pair is U6-A11 |
| **ΔG= -7.80 kca/mol** | | |

**Supplementary Table 4. mFold analysis of MOPV IGR elements**

| IGR-L structural element | δG | Information |
|---|---|---|
| External loop | 0.00 | 0 ss bases & 1 closing helices |
| Stack | -3.30 | External closing pair is C1-G39 |
| Stack | -3.30 | External closing pair is C2-G38 |
| Stack | -3.30 | External closing pair is C3-G37 |
| Stack | -3.30 | External closing pair is C4-G36 |
| Stack | -2.40 | External closing pair is C5-G35 |
| Stack | -2.40 | External closing pair is G6-C34 |
| Stack | -2.10 | External closing pair is A7-U33 |
| Stack | -2.40 | External closing pair is G8-C32 |
| Stack | -2.20 | External closing pair is A9-U31 |
| Stack | -3.30 | External closing pair is C10-G30 |
| Stack | -3.30 | External closing pair is C11-G29 |
| Stack | -2.10 | External closing pair is C12-G28 |
| Stack | -2.20 | External closing pair is A13-U27 |
| Stack | -3.30 | External closing pair is C14-C26 |
| Stack | -2.40 | External closing pair is C15-G25 |
| Stack | -3.40 | External closing pair is G16-C24 |
| Stack | -3.30 | External closing pair is C17-G23 |
| **Helix** | -48.00 | 18 base pairs |
| Hairpin loop | 5.70 | External closing pair is C18-G22 |
| **ΔG= -42.30 kca/mol** | | |
| **IGR-S structural element** | **δG** | **Information** |
| External loop | -0.30 | 1 ss bases & 1 closing helices |
| Stack | -3.30 | External closing pair is C2-G39 |
| Stack | -3.30 | External closing pair is C3-G38 |
| Stack | -3.30 | External closing pair is C4-G37 |
| Stack | -3.30 | External closing pair is C5-G36 |
| Stack | -2.40 | External closing pair is C6-G35 |
| Stack | -3.30 | External closing pair is G7-C34 |
| Stack | -3.30 | External closing pair is G8-C33 |
| Stack | -3.30 | External closing pair is G9-C32 |
| Stack | -3.30 | External closing pair is G10-C31 |
| Stack | -3.30 | External closing pair is G11-C30 |
| Stack | -2.40 | External closing pair is G12-C29 |
| **Helix** | -34.50 | 12 base pairs |
| Interior loop | -1.00 | External closing pair is A13-U28 |
| Stack | -3.30 | External closing pair is C15-G26 |
| Stack | -3.30 | External closing pair is C16-G25 |
| Stack | -3.30 | External closing pair is C17-G24 |
| Helix | -9.90 | 4 base pairs |
| Hairpin loop | 4.00 | Closing pair is C18-G23 |
| **ΔG= -42.30 kca/mol** | | |

**Supplementary Table 5. mFold analysis of TAR stem-loop structures used here.**

| TAR structural element | δG | Information |
|---|---|---|
| External loop | 0.00 | 0 ss bases & 1 closing helices |
| Stack | -3.30 | External closing pair is G1-C59 |
| Stack | -3.30 | External closing pair is G2- C58 |
| Stack | -2.20 | External closing pair is G3-C57 |
| **Helix** | -8.80 | 4 base pairs |
| **Bulge loop** | 2.90 | External closing pair is U4-C56 |
| Stack | -2.40 | External closing pair is U6-A55 |
| Stack | -2.10 | External closing pair is C7-G54 |
| Stack | -1.50 | External closing pair is U8-G53 |
| Stack | -2.10 | External closing pair is C9-G52 |
| Stack | -1.00 | External closing pair is U10-A51 |
| Stack | -2.10 | External closing pair is G11-U50 |
| Stack | -2.20 | External closing pair is G12-C49 |
| Stack | -0.90 | External closing pair is U13-A48 |
| Stack | -1.30 | External closing pair is U14-A47 |
| Stack | -2.10 | External closing pair is A15-U46 |
| **Helix** | -17.70 | 11 base pairs |
| **Bulge loop** | 0.40 | External closing pair is G16-C45 |
| Stack | -3.30 | External closing pair is C18-G44 |
| Stack | -2.10 | External closing pair is C19-G43 |
| Stack | -2.10 | External closing pair is A20-U42 |
| Stack | -2.40 | External closing pair is G21-C41 |
| **Helix** | -9.90 | 5 base pairs |
| **Bulge loop** | 3.70 | External closing pair is A22-U40 |
| Stack | -2.40 | External closing pair is G26-C39 |
| Stack | -2.10 | External closing pair is A27-U38 |
| Stack | -3.40 | External closing pair is G28-C37 |
| **Helix** | -7.90 | 4 base pairs |
| Hairpin loop | 4.40 | Closing pair is C29-G36 |
| **ΔG= -32.90 kca/mol** | | |

**Supplementary Table 5, continued. mFold analysis of TAR stem-loop structures used here.**

| TAR-5SL structural element | δG | Information |
|---|---|---|
| External loop | 0.00 | 0 ss bases & 1 closing helix |
| Stack | -3.40 | External closing pair is G1-C27 |
| Stack | -2.10 | External closing pair is C2- G26 |
| Stack | -2.10 | External closing pair is A3-U25 |
| Stack | -2.40 | External closing pair is G4-C24 |
| **Helix** | -10.00 | 5 base pairs |
| Bulge loop | -3.70 | External closing pair is A5-U23 |
| Stack | -2.40 | External closing pair is G9-C22 |
| Stack | -2.10 | External closing pair is A10-U21 |
| Stack | -3.40 | External closing pair is G11-C20 |
| **Helix** | -7.90 | External closing pair is 4 base pairs |
| Hairpin loop | 4.40 | External closing pair is C12-G19 |
| **ΔG= -9.80 kcal/mol** | | |
| **TAR-9SL structural element** | **δG** | **Information** |
| External loop | 0.00 | 0 ss bases & 1 closing helix |
| Stack | -3.30 | External closing pair is G1-C35 |
| Stack | -3.30 | External closing pair is G2- C34 |
| Stack | -3.30 | External closing pair is G3-C33 |
| Stack | -3.40 | External closing pair is G4-C32 |
| Stack | -3.30 | External closing pair is C5-G31 |
| Stack | -2.10 | External closing pair is C6-G30 |
| Stack | -2.10 | External closing pair is A7-U29 |
| Stack | -2.40 | External closing pair is G8-C29 |
| **Helix** | -23.20 | 9 base pairs |
| Bulge loop | -3.70 | External closing pair is A9-U27 |
| Stack | -2.40 | External closing pair is G13-C26 |
| Stack | -2.10 | External closing pair is A14-U25 |
| Stack | -3.40 | External closing pair is G15-C24 |
| **Helix** | -7.90 | External closing pair is 4 base pairs |
| Hairpin loop | 4.40 | External closing pair is C16-G23 |
| **ΔG= -23.00 kca/mol** | | |
